# Identifying multimodal molecular programs with mTopic

**DOI:** 10.64898/2025.12.21.695756

**Authors:** Piotr Rutkowski, Natalia Ochocka, Damian Panas, Marcin Tabaka

## Abstract

The simultaneous profiling of diverse molecular modalities offers unprecedented insight into complex biological processes, yet it poses significant computational challenges. Here, we introduce mTopic, a generalizable topic modeling framework for analysis of unlimited types of modalities across both single-cell and spatial contexts. mTopic enables identification of coherent multimodal molecular programs at enhanced resolution and supports investigation of cross-modality associations.

---

Genome-wide multimodal sequencing enables simultaneous profiling of diverse modalities, including the epigenome, transcriptome, and epitopes in single cells, as well as within a spatial context [1–4]. mTopic performs an unsupervised, hierarchical decomposition of multimodal data, identifying coherent substructures - referred to as topics - along with modality-specific feature contributions (Fig. 1a). Available topic models are limited to unimodal data [5], tailored protocols [6], or unimodal data integrated with spatial information [7, 8]. In contrast, mTopic extends the Gaussian-multinomial latent Dirichlet allocation (GM-LDA) model [9] to generalize across multiple data modalities and incorporate spatial context. It employs variational inference [10] to iteratively optimize a topic-cell and set of feature-topic probability distributions (Methods), offering an efficient alternative to the Gibbs sampling techniques used in previous methods [5, 6]. mTopic is implemented in Python, integrates with widely used single-cell analysis tools in both R and Python, and supports interactive exploratory analysis of results via Dash.

**Fig. 1.**
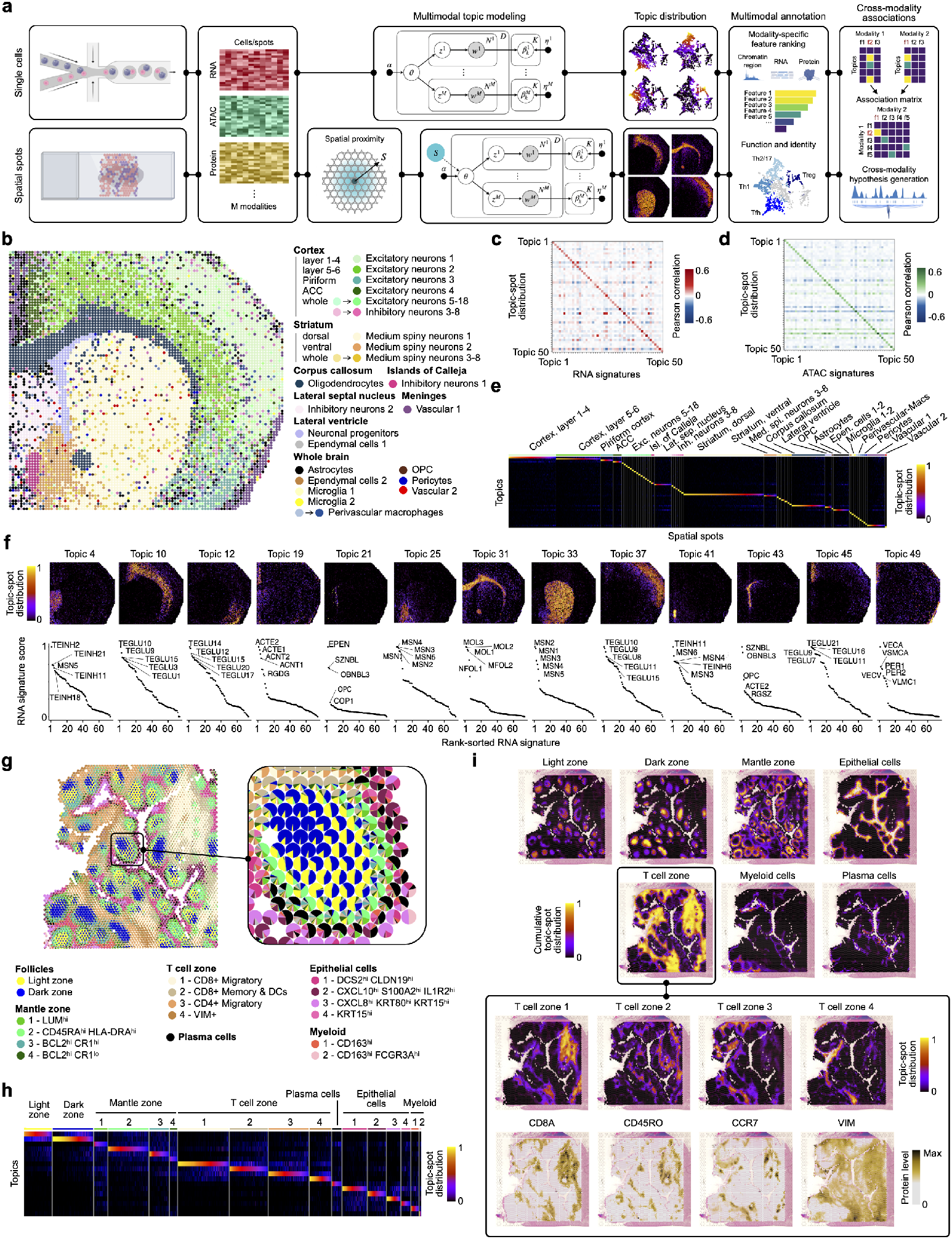
Overview of the mTopic workflow and its application to spatial multimodal data analysis. **a**, Schematic representation of the mTopic workflow. **b**, Cell types identified in the bimodal P22 mouse brain dataset (RNA, ATAC) [4]. **c-d**, Correlation heatmap showing that both RNA and ATAC signatures correlate with the topic-spot distribution (proportion of each topic per spatial spot), indicating consistent topic-specific patterns across modalities. **e**, Heatmap of topic-spot distribution. **f**, Spatial localization of topics across the brain aligns well with the top cell type RNA signature scores calculated per topic using cell type signatures from Adolescent Mouse Brain Atlas [11]. **g**, Cell identities for the Human tonsil spatial dataset [20], pie charts represent topic-spot distribution, each topic has been assigned to one cell identity. **h**, Heatmap of topic-spot distribution. **i**, Spatial cell identity distribution across human tonsil tissue. The upper panel shows the sum of topics for major cell identities. The bottom panel demonstrates the distribution of 4 individual topics of the T-cell zone and their co-localization with the expression of discriminative protein markers.

To illustrate the principles of mTopic, we applied it to spatial epigenome-transcriptome datasets of mouse brain [4]. On the chromatin accessibility and transcriptome (ATAC-RNA) dataset, mTopic correctly identified the spatial distribution of structures that correspond to anatomical regions of the mouse brain (Fig. 1b, Supplementary Fig. 1a, Supplementary Table 1). mTopic results based on 50 multimodal topics show a high degree of concordance between inferred topic-spot distribution and computed RNA signatures (Fig. 1c) or ATAC (Fig. 1d), with topics exhibiting high specificity (Fig. 1e). We annotated multimodal topics based on RNA signatures, leveraging the gene expression values and cell type annotations from the single-cell Adolescent Mouse Brain Atlas [11] (Fig. 1b,f, and Supplementary Table 1). The assigned annotations align well with known cell types and anatomical structures, supporting the robustness and accuracy of mTopic. For example, in cortical layers, we identify Topic 10 enriched in layers 5-6 excitatory neurons (TEGLU10, TEGLU3) and Topic 37 enriched in TEGLU8, specific to layer 4. Other excitatory populations include TEGLU12 and TEGLU15/17 in topic 12, associated with the lateral cortex and piriform area, and Topic 45 (TEGLU21) associated with the anterior cingulate cortex (ACC). Accordingly, topics associated with inhibitory neurons localized in areas rich in GABAergic neurons: Topic 4 in the lateral septal nucleus (TEINH2) and Topic 41 in the islands of Calleja (TEINH11). In the striatum, mTopic correctly identified topics with medium spiny neurons (MSN) signatures: Topic 25 in the ventral striatum (MSN4, D1 neurons) and Topic 33 in the dorsal striatum (MSN2, D2 neurons).

Topic 31 (corpus callosum) is associated with both mature (MOL1-3) and newly formed oligodendrocytes (NFOL), reflecting myelination of neuronal tracts. Topic 19 distributed throughout the brain shows strong astrocyte specificity (ACTE1, ACNT1/2). In addition to major brain structures, mTopic discriminated less abundant populations: Topic 43 enriched in progenitor cells (SZNBL - neuronal intermediate progenitors, OPC - oligodendrocyte precursor cells) is localized within the neurogenic niche next to the lateral ventricle, Topic 21 demarcates ependymal cell layer (EPEN) lining the ventricle; and Topic 49 (meninges) showing vascular enrichment: endothelial cells (VECA/VECV), smooth muscle (VSMCA), pericytes (PER1/2), and vascular leptomeningeal cells (VLMCs). Overall, mTopic robustly captured complex brain architecture, demonstrating its power to dissect cellular diversity within spatial context. The major brain regions have also been recapitulated by spatial distribution patterns of multi-modal topics from H3K4me3, H3K27me3, and H3K27ac histone modification - transcriptome datasets (Extended Data Fig. 1a-c, Supplementary Fig. 1b, 2a-b, and Supplementary Table 1). Moreover, we observed high concordance of RNA rankings between ATAC-RNA and histone modification-RNA corresponding topics, for example RNA probabilities in topics assigned to lateral ventricle or corpus callosum were correlated (Kendall’s *τ* coefficient (0.66, 0.87), *P <* 2.2 × 10^−16^) with similar RNA rankings (Rank Biased Overlap, RBO, p-value *P <* 10^−5^) (Extended Data Fig. 1d). Overall, mTopic more accurately preserved local cell neighborhoods (Extended Data Fig. 1e) and produced feature signatures that were more complex and specific as compared to state-of-the-art multimodal methods [12–16] (Extended Data Fig. 1f). mTopic is scalable, enabling analysis of datasets with over one million cells (Extended Data Fig. 1g).

Next, we assigned chromatin regulatory elements to genes using GREAT [17] to compare chromatin and RNA-topic distributions (Extended Data Fig. 2a-b). H3K27me3, a repressive histone mark, shows minimal correlation at promoters (*r* = 0.24) between feature-topic distributions across all topics, compared to ATAC, H3K4me3, and H3K27ac (Pearson correlation coefficient, *r* =0.62, 0.63, 0.70, respectively), with an even lower correlation at distal elements (*r* = 0.13). We hypothesized that the mutually exclusive H3K27me3 and RNA modalities would exhibit greater signal diversity and reveal finer substructures in the mouse brain compared to RNA-correlated histone modifications. Indeed, mTopic found more dominant topics in H3K27me3-RNA dataset (Extended Data Fig. 2c). Next, we computed RBO values for RNA rankings between the ATAC-RNA and H3K27me3-RNA datasets, linking topics based on their overlap rankings. This revealed a pronounced separation of cortical layers in the H3K27me3-RNA dataset (Extended Data Fig. 2d–e). Among the top-ranked RNA signatures are genes specific to cells forming contiguous cortical layers (Extended Data Fig. 2f): topic 19 — TEGLU2 (layer 6b); topic 2 - TEGLU3 (layer 6) and TEGLU10 (layer 5); topic 46 - TEGLU4 (layer 5); topic 48 - TEGLU8 (layer 4); topic 32 - TEGLU7 (layer 2/3). In addition, ATAC-RNA topic 12 has been split into: topic 1 - TEGLU12 (lateral cortex layer 6); and topic 33 - TEGLU15, 17 (piriform). These results highlight the significant role of H3K27me3 in cortical layer patterning, as further supported by differential analysis (Extended Data Fig. 2g). Moreover, in the ATAC-RNA dataset, we identified topic 31 that co-localized closely with the corpus callosum. This region in the H3K27me3-RNA dataset was divided into two topics (Extended Data Fig. 2h). Topic 14, which encloses the entire corpus callosum structure and shows enrichment of mature oligodendrocytes signature (MOL1-3), and topic 45, which is localized centrally and shows an enriched presence of newly formed oligodendrocytes (NFOL1). This aligns with studies showing continuous renewal of oligodendrocytes in the adult corpus callosum [18] and remyelination occurring predominantly in the central region of the corpus callosum [19].

Next, we applied mTopic to the spatial mutimodal RNA and protein epitope dataset obtained from human tonsil with Visium CytAssist [20]. mTopic identified extrafollicular and follicular zones and discriminated specialized microenvironments within follicular germinal centers (GC), extrafollicular T-cell zones, and epithelium (Fig. 1g, Supplementary Fig. 3a, Supplementary Table 1). The mTopic-generated topic-cell distributions showed high specificity (Fig. 1h) and corresponded well with known tonsil architecture, allowing us to define six major microenvironments: follicles, mantle zone, T-cell zone, epithelial cells, myeloid cells, and plasma cells (Fig. 1i). Within these microenvironments, we further identified 17 distinct functional subsets (Fig. 1h, Supplementary Fig. 3a). Within follicles, the Light Zone (LZ) was marked by high expression of LMO2, a gene upregulated in GC B cells [21], and a cytokine-receptor pair CXCL13 and CXCR5 that directs T follicular helper (TFH) cells to the follicular regions [22] (Extended Data Fig. 3a-b). Spots enriched for the Dark Zone (DZ) signature exhibited elevated expression of proliferation markers MKI67 and TOP2A, along with PCNA protein, which demarcate the DZ within tonsil [23]. In the T-cell zone, we discriminate 4 functional topics (Fig. 1h). T-zone-1 enriched in CD8^+^ T cells end T-zone-3 enriched in CD4^+^ T cells. Both are characterized by the expression of cytokines CCL19 and CCL21 and the homing receptor CCR7, which guide immune cells to secondary lymphoid tissues. T-zone-2 emerged as a potential niche for CD8^+^ memory T cells and dendritic cells (DCs), showing the highest expression of CD45RO and CD11c proteins, alongside elevated CXCL10 expression—a cytokine that facilitates interactions between CD8^+^ T cells and DCs [24]. Whereas, T-zone-4 showed a moderate level of both CD4 and CD8 proteins, upregulation of plasma cells marker MZB1, and high CD27 levels that are shown to be expressed in the extrafollicular zone by T, B, and plasma cells [25]. The identified topics align well with findings from a recent human tonsil atlas, which explored cellular heterogeneity in human tonsils using 5 modalities [26]. While this study annotated 121 distinct cell types based on the single-cell datasets, its spatial transcriptomics analysis revealed only a few broad clusters. In contrast, mTopic achieved high-resolution discrimination of cellular signatures without the need for accompanying single-cell data. In addition to cell identity annotation, we investigated the composition and local context of the 38 identified follicles (Extended Data Fig. 3c). We computed radial profiles by measuring the orientation of cell populations relative to island centroids and clustered the resulting profiles, which revealed 6 follicle groups (Extended Data Fig. 3d-g). The identified clusters might reflect distinct stages of B cell differentiation within germinal centers. For instance, Cluster 2 was dominated by DZ, whereas Cluster 3 showed a strong enrichment of plasma cells, indicative of terminal differentiation and GC exit. These spatial and cellular patterns align with the architectural dynamics of germinal centers, where evolving proportions of LZ, DZ, and plasma cells reflect the maturation trajectory of B cells [27].

Finally, we sought to validate the mTopic model on the droplet-based single-cell PBMC DOGMAseq dataset [3] comprising three modalities: ATAC, RNA, and surface protein. All three modalities were employed for mTopic, and the resulting topic-cell distributions were used for dimensionality reduction (Fig. 2a, Supplementary Fig. 3b). mTopic enabled a clear separation of cell identities (Fig. 2b). Our analysis successfully recapitulated the major cell types identified in the original study [3] - CD8, CD4 T cells, B cells, DCs, Natural Killer (NK) cells, Mucosal-associated invariant T (MAIT), *γδ*T, and Hematopoietic Stem (HSC) cells. mTopic showed high performance in preserving local cell neighborhoods (Extended Data Fig. 4a) and generating high-specificity feature signatures (Extended Data Fig. 4b). The annotated cell types corresponded closely with protein marker levels, with clear separation of naïve and effector fractions based on CD45RA and CD45RO expression (Extended Data Fig. 4c-d). Notably, mTopic enabled distinguishing finer CD4 T-cell subpopulations, such as Th1, Th2/17, Tfh, and Tregs characterized by increased expression of master transcription factors (BCL6, TBX21, GATA3, RORC and FOXP3) and key protein markers (Extended Data Fig. 4e-f). While transcription factor expression is often difficult to detect due to transient expression, we observed BCL6 enrichment in Tfh cells, albeit it was also found in other CD4 subsets. However, the Tfh-associated protein marker CD278 (ICOS) showed specific enrichment in the Tfh cells. Additionally, we identified a rare CD45RA/CD45RO double-positive Treg population, characterized by FOXP3 expression and a medium level of CD25 (Extended Data Fig. 4e-f), which was shown to be present in adult blood [28]. This highlights the importance and advantage of mTopic in integrated analysis of multiple modalities for accurate cell state identification.

**Fig. 2.**
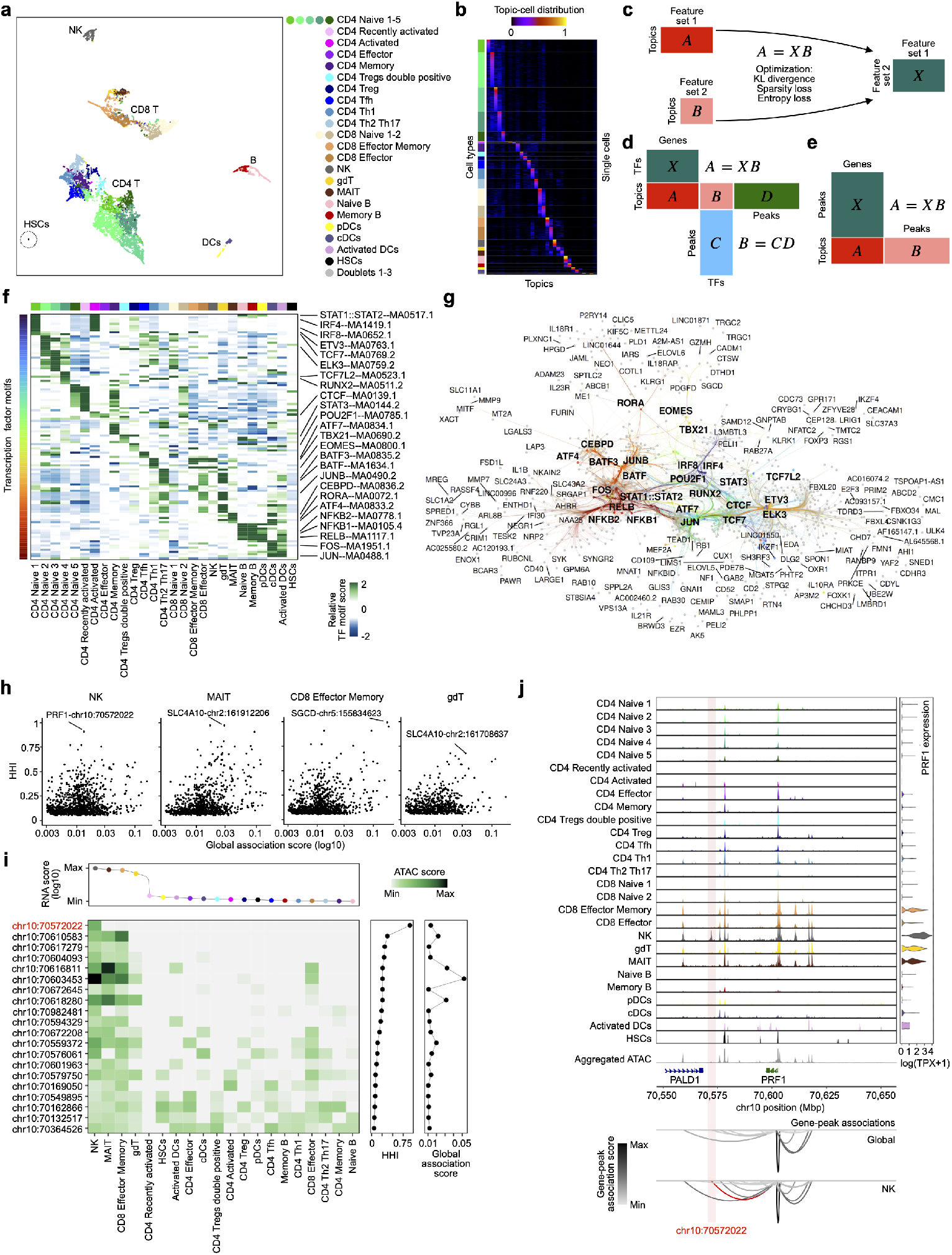
mTopic unravels multimodal molecular programs in peripheral blood mononuclear cells. **a**, UMAP visualization of single cells from topic-cell distributions and assigned cell identities in the PBMC dataset [3] **b**, Heatmap of topic-cell distribution. **c**, Schematic depicting an analytical framework for discovering cross-modality feature relationships from multimodal topic modeling. **d-e**, Matrix block diagrams depicting inference of TF-gene (d) and peak-gene (e) associations. **f**, Relative TF motif scores for each topic. TFs are restricted to FDR≤ 0.01 over the background peaks. **g**, Visualization of the GRN for TFs from (f). TF target nodes are restricted to genes predicted by the method in (d). A subset of nodes is labeled with gene symbols for clarity. **h**, Peak-gene associations in the subnetwork of TBX21, EOMES, and RORA. HHI indicates the specificity of peak-gene association. Top HHI associations are labelled per cell type. **i**, Single-gene characteristics of regulatory landscape inferred by mTopic for PRF1. **j**, Chromatin-accessibility and expression profiles across cell types for PRF1 gene. Loops denote gene-peak associations determined using the global association method in (e) or ATAC scores for individual cell identity. Loop height represents the gene-peak association score.

To uncover cross-modality associations, we developed a framework that links feature sets via their feature-topic distributions (Fig. 2c). It infers an association matrix that minimizes the divergence between the feature-topic distributions of the two modalities. While the framework is broadly applicable to study diverse associations, we applied it here to identify (i) transcription factor (TF)–target gene relationships within gene regulatory networks (GRNs) (Fig. 2d), and (ii) cis peak–gene associations (Fig. 2e). First, we restricted TFs to those with significantly enriched motifs (FDR ≤ 0.01) in the ATAC signatures (Fig. 2f). Enrichment analysis accurately recovered canonical TFs in distinct cell types, e.g., STAT1 and STAT2 in activated CD4 T cells, TBX21 in Th1 cells, RORA in MAIT cells, and NFKB1/NFKB2 in B cells (Fig. 2f). Interestingly, CD4 naive T cells showed enrichment of activation-associated TF motifs in open chromatin regions, despite elevated level of CD45RA - a marker of naive T cells (Extended Data Fig. 4c-d), suggesting early chromatin remodeling in those cells. Then, expressed TFs with enriched motifs were used to learn GRN (Methods) and visualized as a graph (Fig. 2g). For example, CEBPD, a key regulator in myeloid immunity, was enriched in DCs, with MMP9 identified as a direct target, consistent with its known promoter binding and transcriptional activation [29]. In B cells, mTopic identified NFKB1 and RELB that are critical for activation and differentiation of B cells, and their canonical targets CCR7 and CD40, in line with their roles in promoting BCR pathway gene expression and B cell survival [30].

The cross-modality association framework, when applied to infer cis peak–gene links, identified 146, 040 pairs (Extended Data Fig. 5a) across all topics. These links, together with ATAC fragment counts in peaks, accurately predicted RNA transcript levels (Extended Data Fig. 5b). However, predictive performance was reduced in low-abundance cell populations (*<*50 cells) (Extended Data Fig. 5c).

To illustrate the utility of mTopic in dissecting GRNs and chromatin regulatory elements, we focused on a subnetwork comprising TFs - TBX21, EOMES, and RORA (Extended Data Fig. 5d–e), whose expression is enriched in NK cells, CD8^+^ T effector, effector memory, *γδ*T, and MAIT cells (Extended Data Fig. 5f). NK cells exhibited the highest gene target signature scores for TBX21 and EOMES, whereas MAIT cells were enriched for RORA target genes (Extended Data Fig. 5g). Subnetwork target genes were associated with cell killing and leukocyte-mediated cytotoxicity (Extended Data Fig. 5h). Direct targets included PRF1 and granzyme genes (GZMA, GZMB, GZMH), consistent with their cooperative activation [31]. Cis peak–gene links inferred from the cross-modality framework represent global associations across all topics, in contrast to ATAC rankings, which order cell state–specific associations. mTopic further enabled dissection of gene regulatory elements of target genes (Fig. 2h) and within an individual gene across cell identities (Fig. 2i). To quantify peak dominance across topics, we used the Herfindahl–Hirschman Index (HHI), which captures the extent to which a feature is concentrated in a small number of topics and reaches 1 under perfect specificity. For example, among the target genes, peak chr10:70572022 linked to PRF1 is the top unique regulatory element to NK cells (Fig. 2h). PRF1 shows the highest expression in NK cells, and chr10:70572022 is the sole NK-specific peak among its regulatory elements (Fig. 2i–j). This highlights that mTopic can effectively dissect the landscape of peak–gene associations in cis, enabling the identification of global and specific regulatory elements and offering a framework to explore the regulatory architecture underlying gene expression diversity across cell identities.

Our results demonstrate that mTopic is a scalable and generalizable framework for analyzing large-scale single-cell and spatial multimodal datasets. As multimodal cell atlases increasingly incorporate diverse molecular layers at single-cell and spatial resolution, mTopic is well positioned to uncover the spectrum of cell identities and their underlying multimodal molecular programs. Furthermore, the cross-modality association framework enables genome-wide identification of feature–feature relationships, which can support the generation of hypotheses and target prioritization in perturbation studies, thereby facilitating biological discovery.

## Methods

### Bayesian inference

The inference problem aims to find the conditional distribution *p*(***z***|***x***), where ***z*** = *z*_1:*m*_ is a set of latent variables and ***x*** = *x*_1:*n*_ is a set of observations. Due to Bayes’ theorem, *p*(***z*** |***x***) is a joint distribution *p*(***z, x***) divided by evidence *p*(***x***),

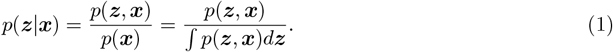

The exact parameter inference is generally impossible, as the evidence integral is usually impossible to compute due to the unavailability of a closed form or the exponential complexity of computations [10]. As a result, approximate algorithms are necessary to estimate *p*(***z***|***x***) parameters, such as Markov chain Monte Carlo [32, 33], Gibbs sampling [33, 34], or variational inference [35].

### Variational inference

Variational inference (VI) is an approach for approximating the posterior distribution using optimization. Let 𝒬 be a family of densities over latent variables ***z***. The idea behind VI is to find an optimal *q*\*(***z***) ∈ 𝒬 that minimizes the Kullback–Leibler (KL) divergence between itself and the posterior distribution *p*(***z***|***x***),

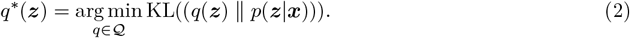

The optimization problem in equation (2) is not yet computable due to dependency on untraceable evidence *p*(***x***),

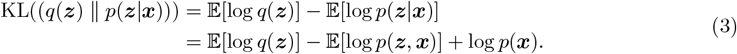

VI omits the evidence *p*(***x***) by using the evidence lower bound (ELBO), defined as negative KL divergence with added evidence,

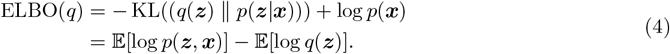

Evidence *p*(***x***) is a constant with respect to *q*(***z***). Hence, minimizing KL divergence is equivalent to maximizing ELBO over the elements of family 𝒬,

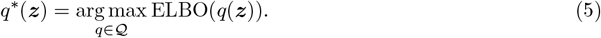

### Mean-field variational family

The quality of a posterior approximation *q*\*(***z***) highly depends on the family of candidate approximations Q. One of the most commonly used families, producing profound distributions while having relatively low complexity, is the mean-field variational family, whose elements are products of variational factors,

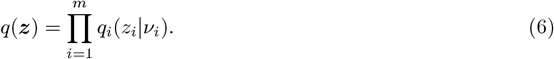

The mean-field variational family assumes that latent variables *z*_*i*_ are mutually independent. Each *z*_*i*_ is governed by its variational factor *q*_*i*_ with variational parameter *v*_*i*_. The mean-field variational family produces optimal variational factors that are proportional to the exponentiated expected log of the complete conditional [10],

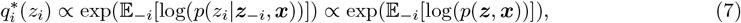

where subscript − *i* refers to all variables except the *i*^th^. The complete conditional of a random variable *z*_*i*_ from the exponential family can be represented as

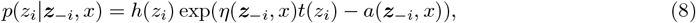

where *h* is a base measure, *η* is a function of the conditioning set, *t* is a sufficient statistic, and *a* is a log normalizer. Using the representation from equation (8), the optimal variational factor from equation (7) simplifies to

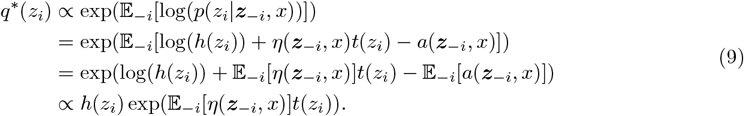

Therefore, *z*_*i*_ and *q*\*(*z*_*i*_) are in the same exponential subfamily. As a result, the update of the *z*_*i*_’s variational parameter is

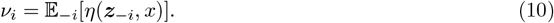

### Choice of distributions

The multimodal topic models introduced in this study use the multinomial (Mult) and Dirichlet (Dir) distributions, both members of the exponential family. This enables application of the parameter update rule defined in equation (10).

The multinomial distribution models discrete outcomes from *N* independent trials over *K* categories, parameterized by a categorical probability vector ***θ*** = (*θ*_1_, …, *θ*_*K*_), where *θ*_*k*_ ≥ 0 and 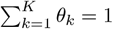. The probability of observing count vector **x** = (*x*_1_, …, *x*_*K*_), such that 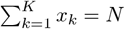, is given by

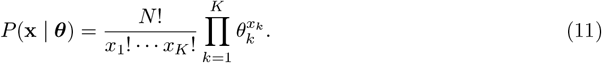

The Dirichlet distribution defines a prior over the *K*-dimensional probability simplex. It is parameterized by a vector of positive concentration parameters ***α*** = (*α*_1_, …, *α*_*K*_) and has the following density

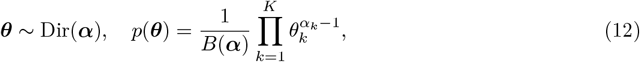

where *B*(***α***) is the multivariate Beta function

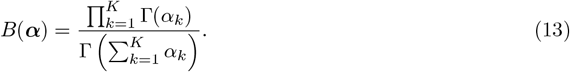

Larger *α*_*k*_ values correspond to higher expected proportions in the *k*-th component. When all *α*_*k*_ are equal, the distribution is symmetric and centered near the uniform distribution.

### Terminology

The terminology used throughout this study is summarized as follows. Multimodal topic modeling produces two primary outputs: (i) a topic–cell distribution, representing topic proportions for each cell, and (ii) a set of feature–topic distributions, comprising one distribution per modality, that represents the contribution of each feature to each topic. Each feature–topic distribution consists of feature probabilities, obtained by normalizing non-negative feature scores generated by the topic modeling framework. Sorting features by their probabilities within a topic yields a feature ranking, with the top-ranked features forming the feature signature of that topic. For a given modality (e.g., RNA), we refer to the corresponding outputs as the RNA–topic distribution, RNA probabilities, RNA scores, RNA ranking, and RNA signature. We use the plural feature–topic distributions to denote the complete set across modalities and the singular feature–topic distribution to refer to a specific distribution for a single modality. In the spatial datasets analyzed here, observations correspond to spots comprising multiple cells; however, the spatial multimodal topic modeling framework is compatible with any spatial resolution.

### Spatial multimodal topic model

The spatial multimodal topic model (sMTM) generalizes the generative model for Latent Dirichlet Allocation (GM-LDA) [9] by extending it to an arbitrary number of data modalities and incorporating spatial context. The model is defined over *D* spatial spots and *M* modalities, where the *m*^th^ modality contains *N* ^*m*^ observed features. Spatial relationships between spots are encoded via a nearest neighbor graph S. Given a fixed number of topics *K*, the model assumes the following generative process,

1. For each of the *M* modalities
  a. For each of the *K* topics
    i. Sample feature distribution 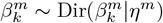
2. For each of the *D* spots
  a. Sample topic-spot distribution *θ*_*dk*_ ∝Dir(*θ*_*dk*_ |*α*)*ξ*_*d*_(𝒮)
  b. For each of the *M* modalities
    i. For each of the *N* ^*m*^ features
      A. Sample topic 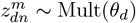
      B. Sample feature 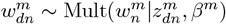,

where *ξ*_*d*_(𝒮) ∈ ℝ^*K*^ is a spatial factor that encodes the topic-spot distribution trends in the spatial neighborhood of spot *d*.

The model specifies the following joint distribution,

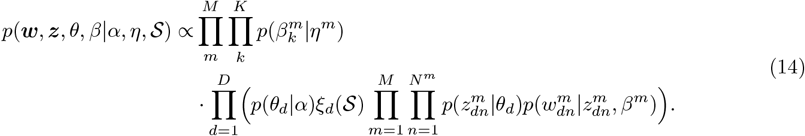

Exact parameter inference for sMTM is intractable. To find the approximated posterior, we employ VI and the mean-field variational family that produces the following factorized distribution of latent variables,

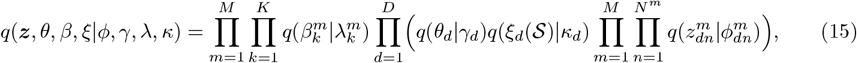

where *ϕ, γ, λ*, and *κ* are variational parameters, each appropriate to its respective latent variable. Following the VI approximation procedure from equation (5), we maximize ELBO using the complete conditional *p*(***w, z***, *θ, β*|*α, η*, S) and the variational function *q*(***z***, *θ, β, ξ*|*ϕ, γ, λ, κ*).

### Update of topic assignment parameters

The topic assignment for the *n*^th^ feature in modality *m* and spot *d*, denoted 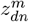, is a multinomial distribution that has the following complete conditional,

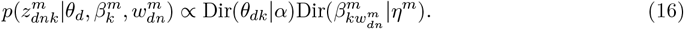

After applying parameter update rule from equation (10), we get the update for the variational topic assignment 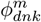,

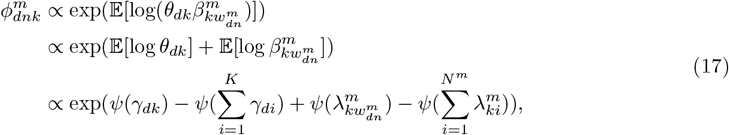

where *γ* are variational parameters of topic-spot distribution *θ, λ* are variational parameters of feature distributions *β*, and *ψ* is a digamma function, defined as the logarithmic derivative of the gamma function,

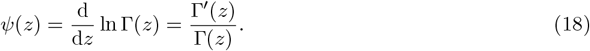

### Update of topic-spot distribution parameters

The topic-spot distribution for spot *d*, denoted *θ*_*d*_, has the following complete conditional,

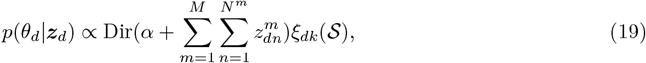

where *ξ*_*dk*_(𝒮) is a spatial scaling factor for spot *d* and topic *k. ξ*_*d*_(𝒮) promotes spatial proximity and the cohesion of topic-spot distributions among neighboring spots. Coupling multiple nearest neighbor topic-spot distributions makes inference a challenge. Inspired by [9, 36], we introduce empirical topic-spot distributions 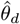. 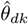can be viewed as a pseudocount that represents the number of times a topic *k* was assigned in spot *d*. During VI, 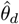 is computed using variational topic assignments *ϕ*,

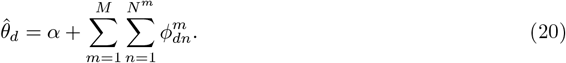

Spatial factors *ξ*_*d*_(𝒮) are computed on empirical topic-spot distributions 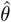 across spatial connections from 𝒮. The impact of a neighboring spot *d*_n_ on the spatial factor *ξ_d_*(𝒮) of spot *d* is positively correlated to Gaussian-scaled spatial proximity and cosine similarity between empirical topic-spot distributions 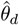 and 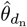. The variational parameter *κ*_*dk*_ of the spatial factor *ξ*_*dk*_(𝒮) is computed as follows. 𝒩 denotes a Gaussian distribution density function with mean *µ* = 0 and standard deviation 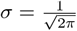. To fit this distribution, spatial coordinates *X*^spatial^ are scaled to be in the range [0, 1]. 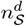 is the number of spatial connections in 𝒮 between spot *d* and other spots,

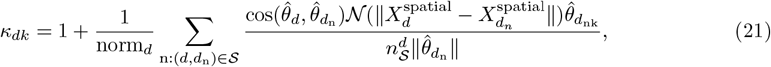

Where

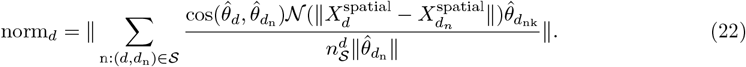

*ξ*_*dk*_(𝒮) is a constant with respect to *θ*_*d*_. Equation (10) and the fact that the expectation of an indicator is its probability give the formula for the update of variational topic-spot distribution parameter *γ*,

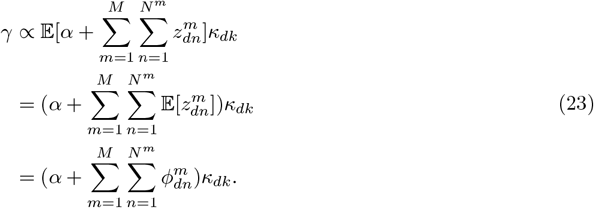

### Update of feature distribution parameters

The feature distribution for topic *k* and modality *m*, denoted 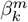, is Dirichlet with the following complete conditional,

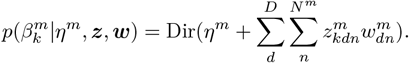

Following Equation (10) and the fact that the expectation of an indicator is its probability, we get the variational feature distribution parameter update,

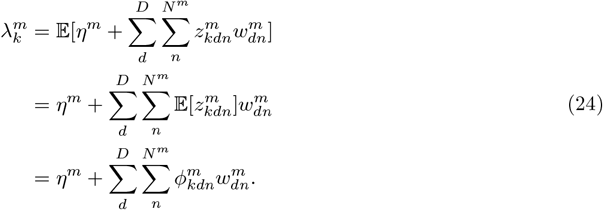

### Training procedure

When training sMTM, we employ coordinate ascent variational inference (CAVI) [37] algorithm to find the optimal approximation *q*\* using parameter updates from equations (17), (23), and (24). Topic assignments and topic-spot distributions are updated iteratively until converged. To omit the exponential impact of the iterative scaling with spatial factors *ξ*(𝒮), we apply spatial factors after each iteration of the topic-spot distributions converges. The training procedure is summarized in Algorithm 1.

### Multimodal topic model

The multimodal topic model (MTM) adapts the spatial multimodal topic model (sMTM) for non-spatial data. The *K*-topic MTM model omits spatial structure and assumes the following generative process, conditioned on *D* cells and *M* modalities, where the *m*^th^ modality contains *N* ^*m*^ observed features,

1. For each of the *M* modalities
  a. For each of the *K* topics
    i. Sample feature distribution 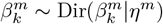
2. For each of the *D* cells:
  a. Sample topic-spot distribution *θ*_*d*_ ~ Dir(*θ*_*d*_|*α*)
  b. For each of the *M* modalities
    i. For each of the *N* ^*m*^ features
      A. Sample topic 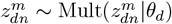
      B. Sample feature 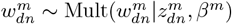.

#### Algorithm 1

Training procedure of sMTM using CAVI

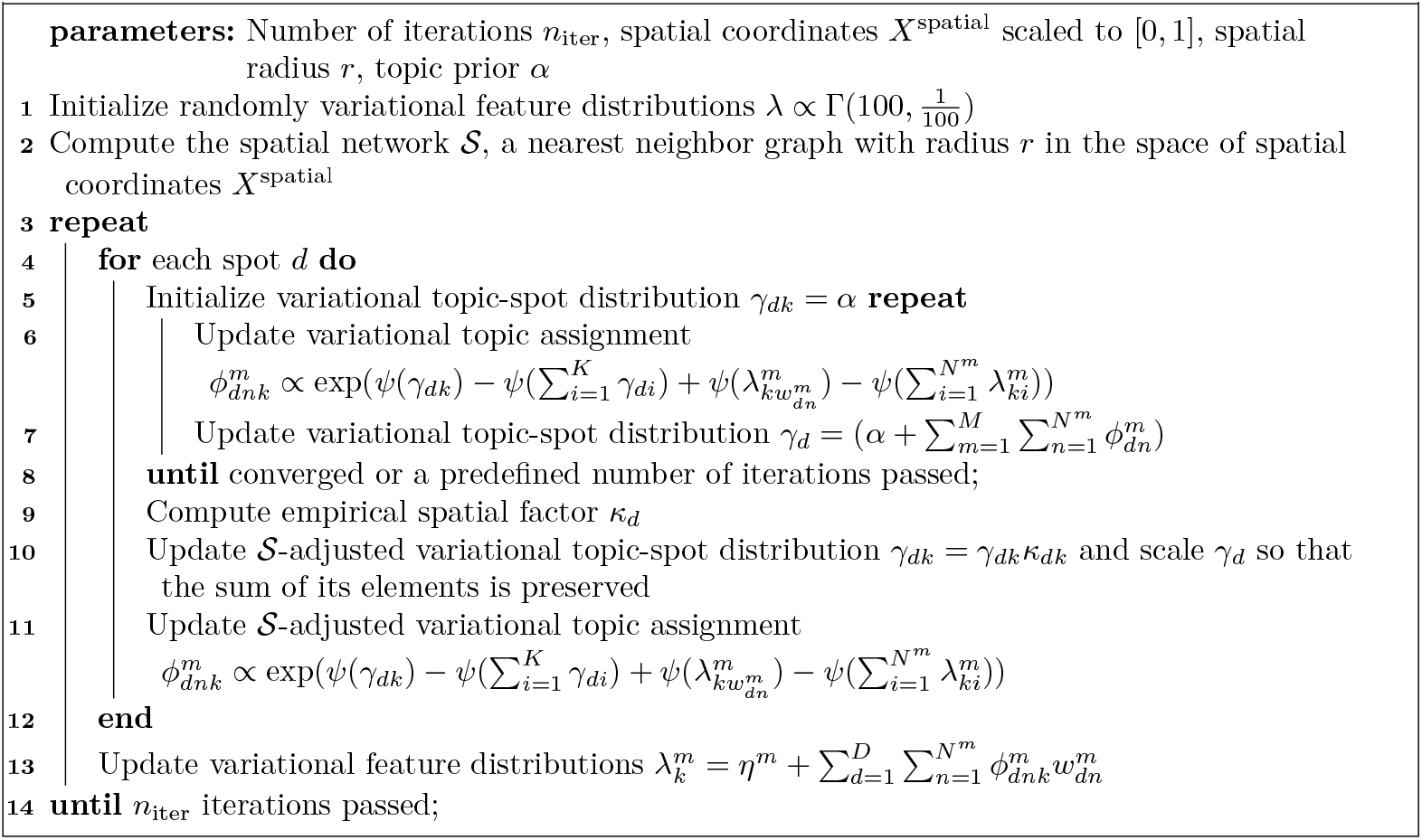

The model specifies the following joint distribution,

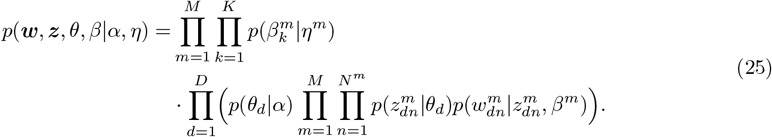

The exact inference for MTM is intractable. Analogously to sMTM, we employ the mean-field VI to find the approximated posterior of the following factorized distribution of latent variables,

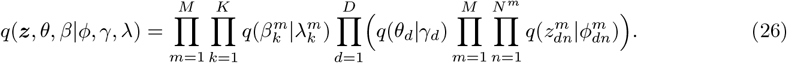

Derivation of parameter updates for the MTM model is the same as for sMTM when omitting spatial factors, resulting in the following updates for the topic assignment variational parameter *ϕ*,

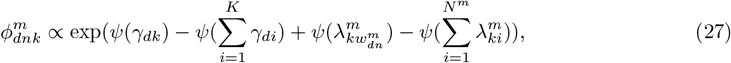

topic-cell distribution variational parameter *γ*,

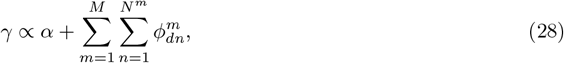

and feature distribution variational parameter *λ*,

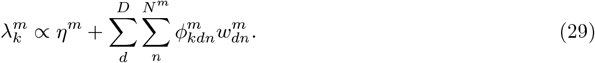

We employ two VI training procedures to find the optimal approximation *q*\*: CAVI [37], summarized in Algorithm 2, and batch stochastic variational inference (SVI) [35], summarized in Algorithm 3. While CAVI updates parameters iteratively using the whole dataset in each iteration, SVI uses random cell batches, making it more suitable for large datasets when speed is required.

#### Algorithm 2

Training procedure of MTM using CAVI

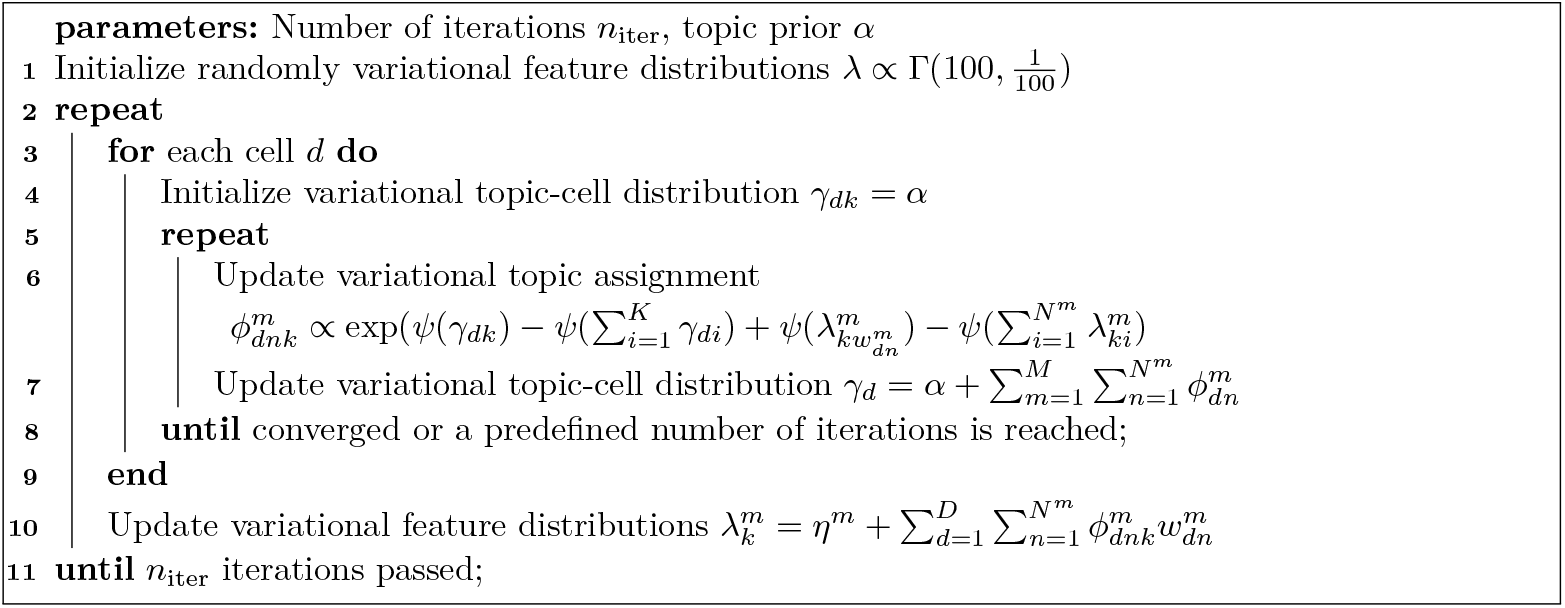

#### Algorithm 3

Training procedure of MTM using SVI

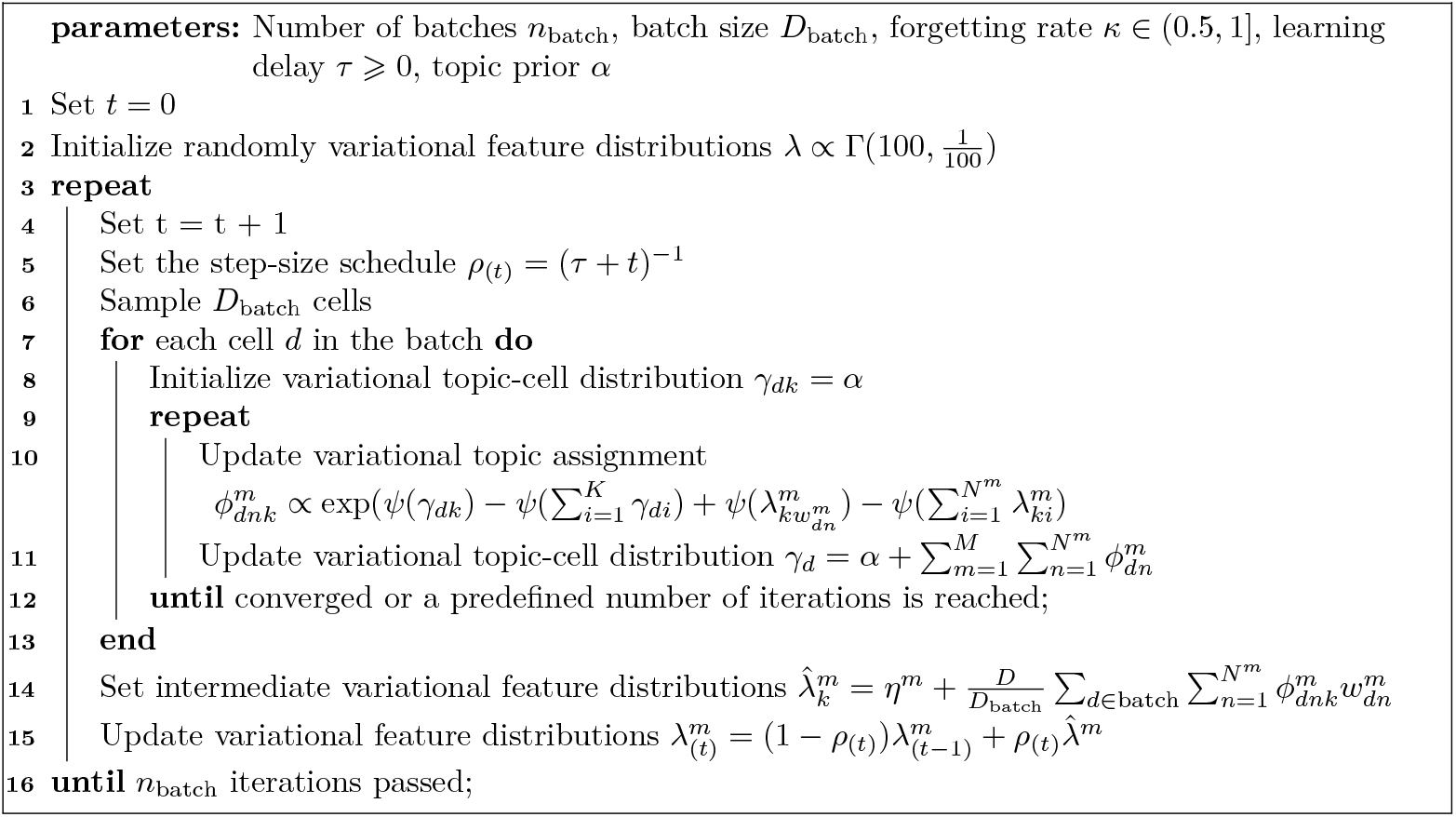

### Initial filtering of pervasive features

Filtering abundant features is essential to ensure that multimodal signatures identified by topic modeling consist of discriminative rather than ubiquitous features. We permute each count matrix along the observation (cell/spot) axis to detect pervasive features likely reflecting technical noise and apply standard preprocessing to the permuted data. RNA, ATAC, and histone modification modalities undergo term frequency–inverse document frequency (TF–IDF) transformation. At the same time, protein epitope data is processed using centered log-ratio (CLR) transformation. Each transformed matrix is then scaled such that the total counts per cell average 10, 000, preserving relative proportions and harmonizing total counts across modalities.

We subsequently train a control MTM model on the permuted data to identify features that dominate feature-topic distributions despite randomized input. Features are ranked by the sum feature–topic scores across all topics, and the inflection point in the sum curve is detected using the Kneedle algorithm [38] to select and exclude pervasive features from downstream analysis.

### Identifying cross-modality feature associations

Interpreting cross-modality feature relationships is a challenge. To address this, we learn associations between two feature sets using topic modeling-derived feature distributions. Let 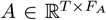 and 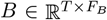 be row-normalized topic–feature distribution matrices with *T* topics and *F*_*A*_ and *F*_*B*_ features, respectively. We define the association matrix between feature sets *A* and *B* as 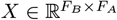 such that

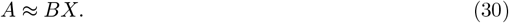

We minimize the discrepancy between the observed matrix *A* and its reconstruction Â = *BX* using a composite loss function,

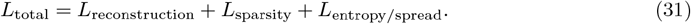

The reconstruction loss *L*_reconstruction_ is a Kullback–Leibler (KL) divergence between the observed distribution *A* and the reconstructed distribution Â,

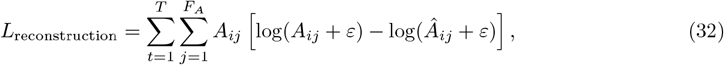

where the constant *ε* ensures numerical stability.

The sparsity loss *L*_sparsity_ encourages sparsity in the association matrix *X*. It combines a fractional norm and logarithmic penalty,

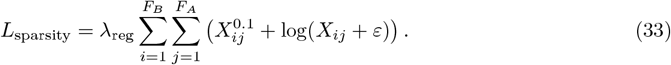

The weighted regularization term *L*_entropy/spread_ balances entropy and spread of values in the association matrix *X*,

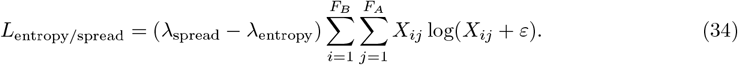

For each feature *a* ∈ {1, …, *F*_*A*_} in *A*, we define a parametric model with learnable logits of length *F*_*B*_, transformed via a softmax function with temperature *τ* to produce a probability distribution over features in *B*,

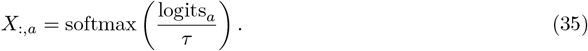

We optimize each *X*_:,*a*_ by minimizing the total loss *L*_total_ using stochastic gradient descent with the Adam optimizer. The model can use custom, feature-feature binary masks that specify for each feature in *A*, which features in *B* should be considered during training.

### Software and dashboard

mTopic was developed in Python (v3.11). It uses the MuData structure [39] to organize multimodal data, enabling seamless integration with existing single-cell analysis workflows. Scalability is achieved through parallelization using joblib (v1.3.2), while cross-modality associations are computed using PyTorch (v2.5.1). mTopic also includes utilities for visualizing model outputs using matplotlib (v3.9.1), and an interactive dashboard built with Dash (v2.15.0) and Plotly (v5.18.0), which enables rapid exploration of mTopic-generated topic-cell and feature-topic distributions. The software, documentation, tutorials, and dashboard setup instructions are available at https://github.com/TabakaLab/mTopic. An R wrapper, tested with R (v4.4.3), is provided for compatibility with Seurat objects. It is based on reticulate (v1.41.0) and was tested with Seurat (v5.2.1). Installation instructions, documentation, and tutorials are available at https://tabalakab.github.io/mTopicR.

### Gene signature analysis

Gene signature scores for the P22 mouse brain datasets were computed using expression matrices and cell type annotations from the Adolescent Mouse Brain Atlas [11], restricted to tissues present in P22 coronal sections at bregma 1. Only atlas clusters with more than 10 cells were included. The RNA signatures of the top 25 genes were used as input to Seurat v5.2.0’s AddModuleScore function (nbin = 20, ctrl = 100), and the resulting gene signature scores were scaled to the (0,1) range.

### Analysis of feature rankings

The correspondence between feature rankings was quantified using Kendall’s *τ* coefficient, computed after intersecting RNA transcripts across epigenomic modalities. Additionally, rank-biased overlap (RBO) [40], as implemented in gespeR (10.18129/B9.bioc.gespeR), was calculated using the full set of RNA transcripts. Statistical significance was assessed by comparing observed values to a null distribution generated from 100,000 random permutations of topic feature rankings.

### Peak-gene cis-association

The peak–gene association framework, applied to the PBMC dataset [3], used a custom mask that linked each gene to peaks located within ±500 kb of its transcription start site (TSS), as defined by the GREAT tool [17]. Before training, we excluded three topics annotated as doublets (topics 9, 19, and 21). The framework was run with default parameters, using the RNA–topic distribution as matrix *A* (27 topics by 5,514 genes shared between the topic model and the GREAT-derived gene list) and the ATAC–topic distribution as matrix *B* (27 topics by 136,915 peaks). In downstream analyses, only peaks with association scores exceeding 1*/N*_peaks_, where *N*_peaks_ is the number of ATAC peaks within the±500 kb window, were retained. These scores are referred to as global peak–gene association scores, in contrast to ATAC topic scores, which capture cell–state–specific associations. Peak–gene associations (as chromatin loops), chromatin accessibility tracks, and gene expression across cell states were visualized using Seurat v5.2.0 and Signac v1.14.0 with custom functions. To identify cell–state–specific peaks within the set of globally associated peaks for each gene, the Herfindahl–Hirschman Index (HHI) was computed from ATAC topic scores across cell states. An HHI of 1*/N*_topics_ corresponds to a uniform distribution of peak scores across topics, whereas an HHI of 1 indicates peak specificity to a single cell state.

### TF motif score

TF motifs with position weight matrices (PWMs) were obtained from the JASPAR2022 core database [41] using TFBSTools [42] in R. Motif matches within 300bp ATAC peaks were identified using the matchMotifs function from the motifmatchr package (doi:10.18129/B9.bioc.motifmatchr) with a stringent p-value cutoff of 10^−5^. Next, for each topic, the top 2,000 peaks ranked by ATAC topic scores were selected. Motif enrichment was assessed by comparing observed motif scores to a null distribution generated from 2 × 10^6^random sequences across genome matched in length to the ATAC peaks. Adjusted p-values were computed using the Benjamini–Hochberg method, and TF motifs with FDR ≤0.01 in at least one topic and TF expressed in *>* 1% of cells were retained for further analysis. Final motif scores were z-score normalized and visualized as a heatmap.

### Gene regulatory network

The TF–gene association framework, applied to the PBMC dataset [3], was run with default parameters. Before training, we excluded three topics annotated as doublets (topics 9, 19, and 21). The TF–topic distribution was obtained by multiplying the ATAC–topic distribution matrix by the TF–ATAC matrix. The RNA–topic distribution was used as matrix *A* (27 topics × 10,000 genes), and the TF–topic distribution as matrix *B* (27 topics × 200 TFs). TF target gene scores, obtained from the cross-modality feature association framework, were z-score normalized, and genes with z-scores *>* 3 were retained for GRN construction. A directed graph was built with TFs as source nodes, target genes as destination nodes, and edge weights corresponding to the z-scores. Gene regulatory networks (GRNs) were constructed using the igraph package [43] and visualized with ggraph [44], employing an edge bundling force layout for improved clarity of network edges. In addition, gene set enrichment analysis (GSEA) [45] was performed on target genes of selected transcription factors.

### Analysis of spatial topic distribution patterns

We identified follicle-associated spots in the human tonsil dataset [20] based on topic profiles and spatial coordinates. First, we selected six topics annotated as follicles (5 and 15) or mantle zones (2, 3, 9, and 14). We classified spots as follicle-enriched if mean activity across selected topics was above the 75th percentile. We constructed a spatial *k*-nearest neighbors graph (*k* = 6) and filtered the graph to retain only follicle-enriched spots. We identified connected components using NetworkX [46] and removed ones with less than 6 spots, creating a set of spatial islands. To capture the local microenvironment around each island, we expanded each island with spots reachable within two graph hops based on shortest-path distances calculated from the spatial graph. For each island, we assigned its centroid to mean coordinates of pre-expansion spots. Next, we binned spots into 16 radial sectors based on their polar angle around the centroid. Within each sector, we averaged topic-spot distributions across all spots, generating radial profiles representing the radial distribution of topics around each island. We computed topic-specific directional vectors for each island by weighing cosine and sine components of radial sectors by radial sector topic scores. Next, we constructed dot product matrices between directional vectors, capturing each island’s relative radial organization of topics. We flattened the upper triangular part of each island’s dot product matrix, stacked them, and generated a Pearson correlation matrix between islands, which we clustered into 6 clusters using hierarchical clustering with average linkage. We averaged dot product matrices among cluster’s spatial spots for each cluster. We computed the first principal component (PC) that captures the significant variation of the spatial organization of topics in each cluster.

### Preprocessing and analysis details of multimodal spatial and single-cell datasets

#### P22 mouse brain: spatial ATAC/H3K4me3/H3K27me3/H3K27ac-RNA

The bimodal P22 mouse brain datasets include ATAC or histone modification (H3K4me3, H3K27me3, and H3K27ac) and RNA modalities, co-profiled on 100× 100 microfluidic chips with 20-*µ*m channels [4]. We generated ATAC and histone modification count matrices with 300bp peaks using ArchR [47], generated spot-peak count matrices, and downsampled them to 10,000 counts per spatial spot. After removing spatial spots outside the tissue, we removed overrepresented features based on cumulative topic-feature parameters from an MTM model trained on permuted count matrices. Knee sensitivities were set to 5 for all modalities. We selected 10,000 highly variable RNA genes for all datasets and 50,000 highly variable ATAC peaks for the ATAC–RNA dataset. The processed datasets contain: (i) ATAC–RNA: 9,215 spatial spots, 50,000 peaks, 10,000 genes; (ii) H3K4me3–RNA: 9,548 spatial spots, 43,355 peaks, 10,000 genes; (iii) H3K27me3–RNA: 9,752 spatial spots, 53,882 peaks, 10,000 genes; (iv) H3K27ac–RNA: 9,370 spatial spots, 45,147 peaks, 10,000 genes. For each of the P22 mouse brain datasets, we trained an sMTM model with 50 topics using VI for 500 iterations.

#### Human tonsil: spatial 10x Visium CytAssist

The FFPE human tonsil dataset includes two modalities, transcriptome and protein epitopes, co-profiled with CytAssist Spatial Gene and Protein Expression and preprocessed with Space Ranger 2.1.0, 10x Genomics (2023, May 15) [20]. After removing spatial spots outside the tissue and proteins highly correlated with isotope control (PDCD1, EPCAM, CD14, CD4, CD163, CR2), we removed overrepresented features based on cumulative topic-feature parameters from an MTM model trained on permuted count matrices. Knee sensitivities were set to 10 for both modalities. We selected 5,000 highly variable transcriptome genes. The processed dataset contains 4,194 spatial spots, 5,000 genes, and 27 proteins. For multimodal analysis with mTopic, we trained an sMTM model with 19 topics using VI for 20 iterations. The feature expression plots have been generated using the Seurat package. The output of sMTM modeling, topic-spot distribution *γ*, has been added to the Seurat object as a separate assay. Heatmaps were drawn with the ‘DoHeatmap’ function on counts for the *γ* assay on scaled RNA and protein assay data.

#### Peripheral blood mononuclear cells: single-cell DOGMA-seq

The peripheral blood mononuclear cells dataset is a control dataset under low-loss lysis (LLL) conditions and includes three modalities, chromatin accessibility, transcriptome, and surface proteins, co-profiled with DOGMA-seq [3]. We generated a chromatin accessibility count matrix with 300bp peaks using ArchR [47], removed isotope control proteins (Rat IgG2b and Rat IgG1k), and peaks detected in less than 30 cells. We kept cells with more than 1,000 gene expression counts, 500 genes expressed, 1,000 chromatin accessibility counts, and 25,000 protein counts. Next, we removed overrepresented features based on cumulative topic-feature parameters from an MTM model trained on permuted count matrices. Knee sensitivities were set to 5 for chromatin accessibility and transcriptome and 0 for proteins. We selected 10,000 highly variable genes. The processed dataset contains 7,349 cells, 136,915 peaks, 10,000 genes, and 208 surface proteins. For multimodal analysis with mTopic, we trained an MTM model with 30 topics using VI for 500 iterations.

### Benchmarks

#### Data preprocessing

Benchmarking was conducted on the PBMC [3], and P22 mouse brain [4] datasets. We evaluated the ability of multimodal methods to preserve local cell neighborhoods using Cobolt [12], Matilda [13], MOFA+ [14], MOJITOO [15], mTopic, and WNN [16]. Additionally, we assessed the specificity of modality-specific feature signatures generated by MOFA+ and mTopic. All methods were applied to identical, filtered count matrices as described in the section *Preprocessing and analysis details of multimodal spatial and single-cell datasets*. Where possible, we fixed the embedding dimensionality across methods for consistency. Exceptions include MOJITOO, which does not allow manual control over the dimensionality of the joint embedding, and WNN, which outputs a neighborhood graph rather than a numerical embedding. To align with the analyses presented in the manuscript, we used 30 embedding dimensions for the PBMC dataset and 50 for the P22 mouse brain datasets. MOJITOO produced embeddings with 42 dimensions for the PBMC dataset, 43 for the P22 ATAC–RNA dataset, 42 for the P22 H3K4me3–RNA dataset, 44 for the P22 H3K27me3–RNA dataset, and 45 for the P22 H3K27ac–RNA dataset.

For Cobolt [12], models were trained on count matrices for 100 epochs, with learning rates of 0.001 and 0.0001 tested; 0.001 yielded the best performance across datasets. Matilda [13] models were trained for 100 epochs, with learning rates of 0.01, 0.001, and 0.0001; 0.001 performed best for P22 datasets, while 0.01 was optimal for the PBMC dataset. We only showed the best scores for Cobolt and Matilda when the benchmarking results were shown. For MOFA+ [14], models were trained on count matrices scaled to unit variance in the slow convergence mode with the maximum number of iterations set to 200. We followed standard Seurat preprocessing for MOJITOO [15] and WNN [16]. ATAC and histone modification modalities (H3K4me3, H3K27me3, H3K27ac) were TF-IDF-transformed and reduced to 50 LSI components (we excluded the first component from model training). RNA count matrices were normalized, standardized, and reduced to 50 PCA components. Protein count matrices were CLR-normalized, standardized, and reduced to 50 PCA components for the PBMC dataset. mTopic models were trained as described in the section *Preprocessing and analysis details of multimodal spatial and single-cell datasets*.

#### Preservation of cell neighborhoods

We assess neighborhood preservation by computing the average overlap of *k*-nearest-neighbor sets between the high-dimensional modality count matrices and their corresponding low-dimensional multimodal embeddings [48], which we call the neighborhood overlap score. We use cosine distance for count matrices and Euclidean distance for embeddings. We compute the cumulative neighborhood overlap score by summing scores computed individually for each modality. We set *k* = 200 for the PBMC dataset [3] and *k* = 20 for the P22 mouse brain datasets [4].

#### Specificity of feature signatures

We assess the specificity of modality-specific feature signatures automatically generated using MOFA+ [14] and mTopic. We compute a Pearson correlation matrix *P* between multimodal embedding components (MOFA+ factors [14] or mTopic topics) and the z-scores of their respective feature signatures (MOFA+ weights [14] or mTopic topic-feature distributions). We compute z-scores on lognormalized count matrices for signatures containing the *n* features with the highest-ranked parameters: *n* = 100 for ATAC and histone modifications (H3K4me3, H3K27me3, H3K27ac), *n* = 20 for RNA, and *n* = 3 for protein epitopes. Next, we evaluate the correlation matrix *P* using a diagonal dominance ratio, defined as:

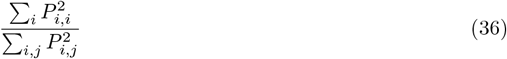

This ratio quantifies the concentration of correlation values along the diagonal of the matrix. A higher score suggests a higher specificity of the generated gene signatures.

#### Evaluating mTopic scalability

To assess the scalability of mTopic, we ran the MTM model on simulated datasets using a 100-core CPU machine running Debian GNU/Linux 12.8. The datasets included 5,000, 10,000, 50,000, 100,000, 500,000, and 1,000,000 cells, each with 2, 3, or 4 modalities comprising 10,000 features per modality. Counts were generated from a Poisson distribution with *λ* = 1 to reflect typical characteristics of count-based single-cell data. All count matrices were constructed with a density level of 5%.

## Code availability

Source code is available under a BSD-Clause 2 license at https://github.com/TabakaLab/mTopic. The documentation and tutorials are available at https://mtopic.readthedocs.io.

## Data availability

Single-cell data analyzed in this study are publicly available from the Gene Expression Omnibus (GEO) under accession numbers: P22 mouse brain datasets (RNA data: [GSE218593](https://www.ncbi.nlm.nih.gov/geo/query/acc.cgi?acc=GSM6753043), specifically GSM6753043 for ATAC-RNA, [GSM6753046](https://www.ncbi.nlm.nih.gov/geo/query/acc.cgi?acc=GSM6753043) for H3K4me3-RNA, GSM6753044 for H3K27me3-RNA, and GSM6753045 for H3K27ac-RNA; ATAC/histone modification data: GSE205055, specifically GSM6758285 for ATAC-RNA, GSM6704980 for H3K4me3-RNA, GSM6704978 for H3K27me3-RNA, and GSM6704979 for H3K27ac-RNA); and human PBMC datasets (GSE166188, specifically GSM5065524 for ATAC, GSM5065525 for RNA, and GSM5065526 for proteins). The human tonsil dataset is available from 10x Genomics (https://www.10xgenomics.com/datasets/gene-protein-expression-library-of-human-tonsil-cytassist-ffpe-2-standard).

## Acknowledgments

We thank members of the Computational Genomics Group for their discussions.P.R. and M.T. are supported by the International Centre for Translational Eye Research (FENG.02.01-IP.05-T005/23) project, which is carried out within the International Research Agendas programme of the Foundation for Polish Science co-financed by the European Union under the European Funds for Smart Economy 2021-2027 (FENG); and grants funded by National Science Center, Poland: the Sonata Bis 12 grant 2022/46/E/NZ2/0037 (M.T.), and Preludium 21 2022/45/N/NZ2/02311 (P.R.).

## Author Contributions

M.T. conceived and supervised the project. P.R. and M.T. designed and developed mTopic. P.R. implemented mTopic and wrote its documentation. M.T., N.O., and P.R. conducted the multimodal data analysis and interpretation of results. D.P. implemented the DASH framework and the R wrapper, and wrote their documentation M.T., N.O., and P.R. wrote the manuscript.

---

**Extended Data Fig. 1.**
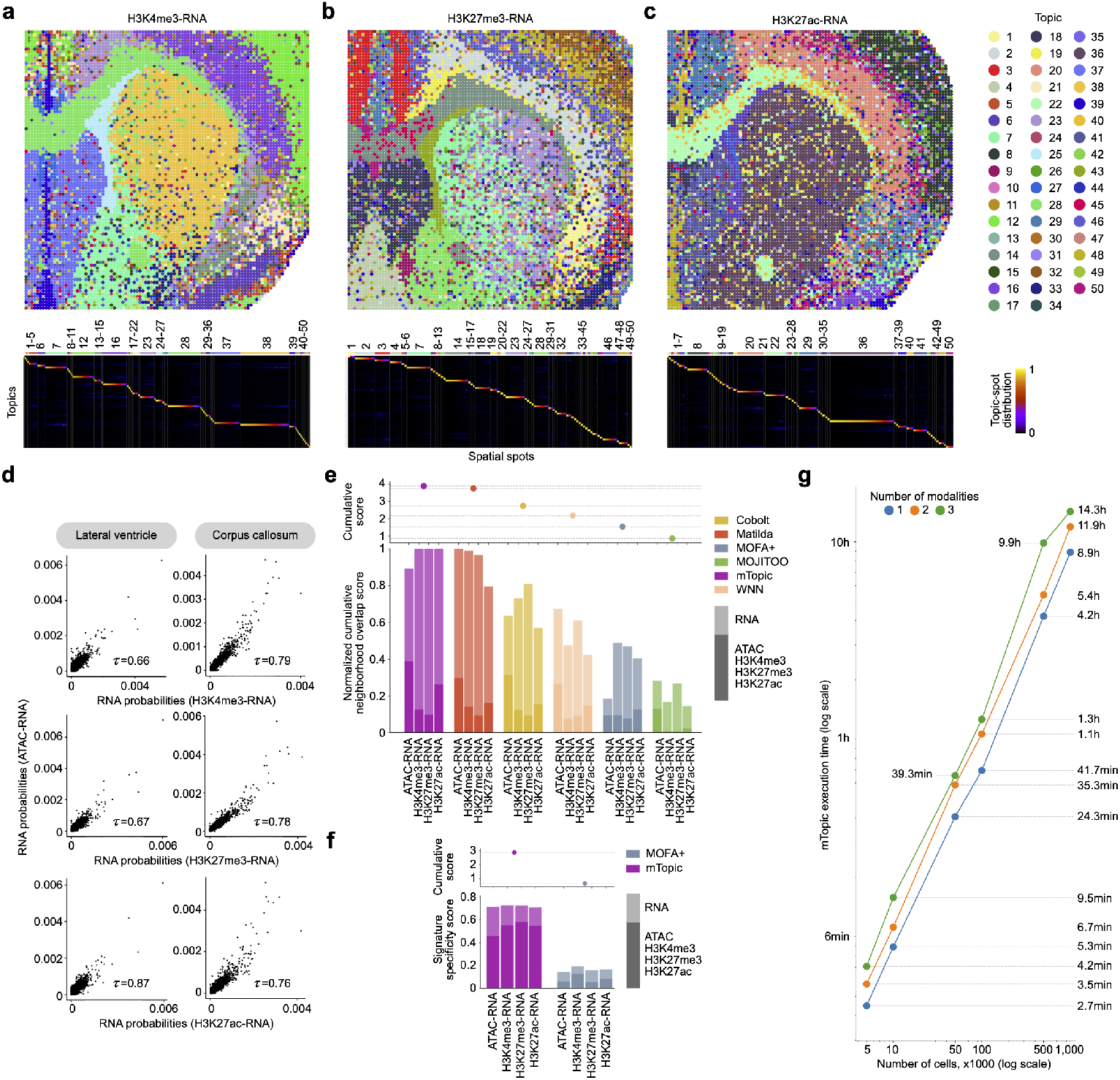
mTopic analysis of histone modification-RNA bimodal datasets from P22 mouse brain. **a-c**, Spatial localization of topics across coronal brain sections (upper panel) and heatmaps of topic-spot distributions (lower panel) for H3K4me3-RNA (a), H3K27me3-RNA (b), and H3K27ac-RNA (c) datasets [4]. **d**, Comparison of RNA probabilities between ATAC-RNA and H3K4me3/K3K27me3/H3K27ac-RNA datasets across lateral ventricle and corpus callosum. **e**, Bench-marking preservation of cell neighborhoods by various multimodal methods applied to chromatin-RNA P22 mouse brain datasets. **f**, Benchmarking specificity of feature signatures between MOFA+ and mTopic. **g**, Benchmarking of execution times of mTopic across various compositions of multimodal datasets.

**Extended Data Fig. 2.**
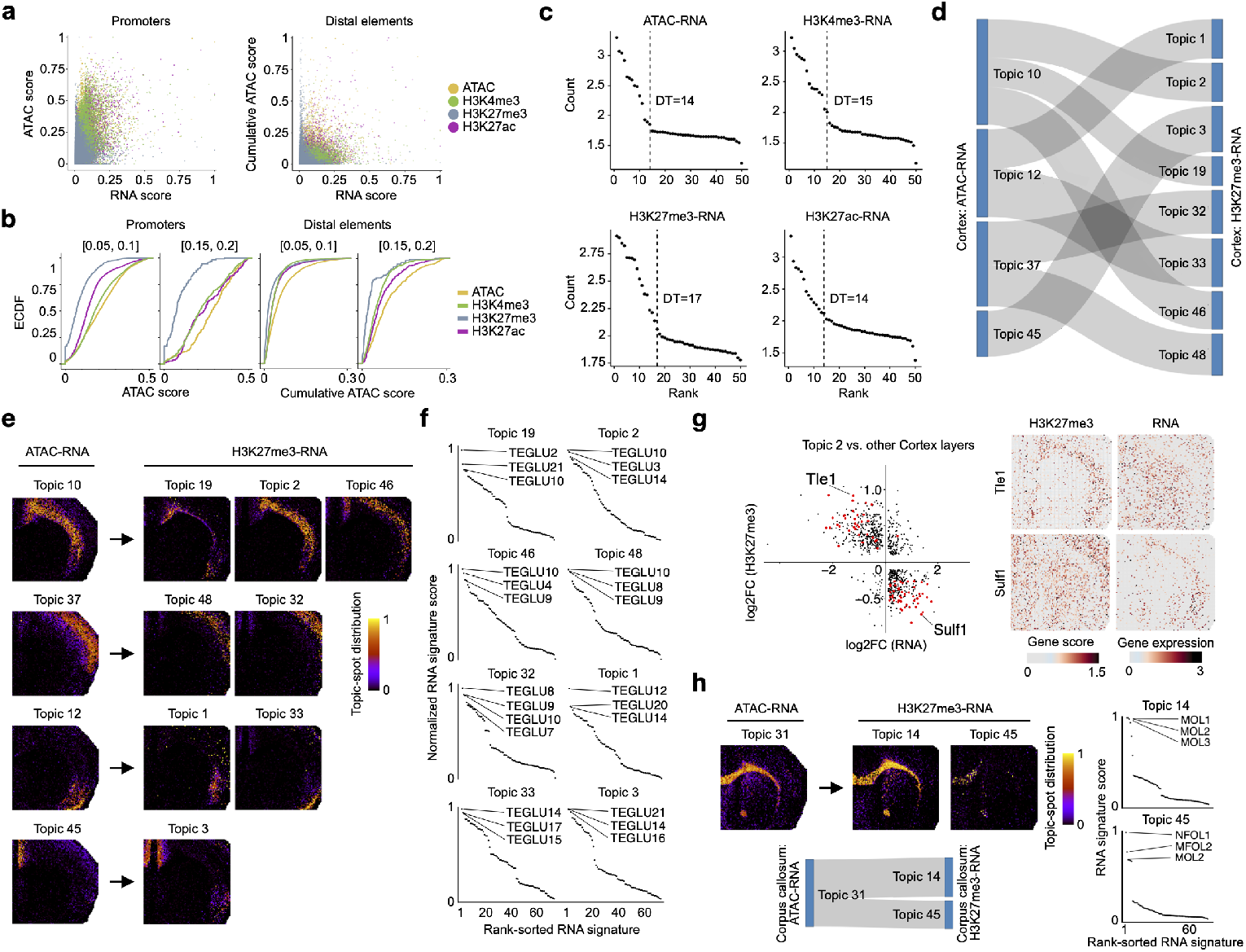
Comparative analysis of bimodal P22 mouse brain datasets. **a**, Correlation between RNA and ATAC scores for promoters and distal elements across all topics. **b**, Empirical cumulative distribution function (ECDF) of ATAC scores in the selected range of RNA scores. **c**, Census of topics across all chromatin-RNA datasets. **d**, Graph representation of relationships among RNA topic rankings between ATAC and H3K27me3 modalities within cortex. Edges represent rank-biased overlap values between topic feature rankings. **e**, Comparison of topic-spot distributions in the cortex between the ATAC-RNA and H3K27me3-RNA datasets. **f**, RNA signature scores calculated per topic in the cortex of the H3K27me3-RNA dataset using cell type signatures from the Adolescent Mouse Brain Atlas [11] **g**, Differential analysis of RNA expression and H3K27me3 histone modification between topic 2 and other cortex layers. **h**, Comparison of topic-spot distributions in the corpus callosum between the ATAC-RNA and H3K27me3-RNA datasets.

**Extended Data Fig. 3.**
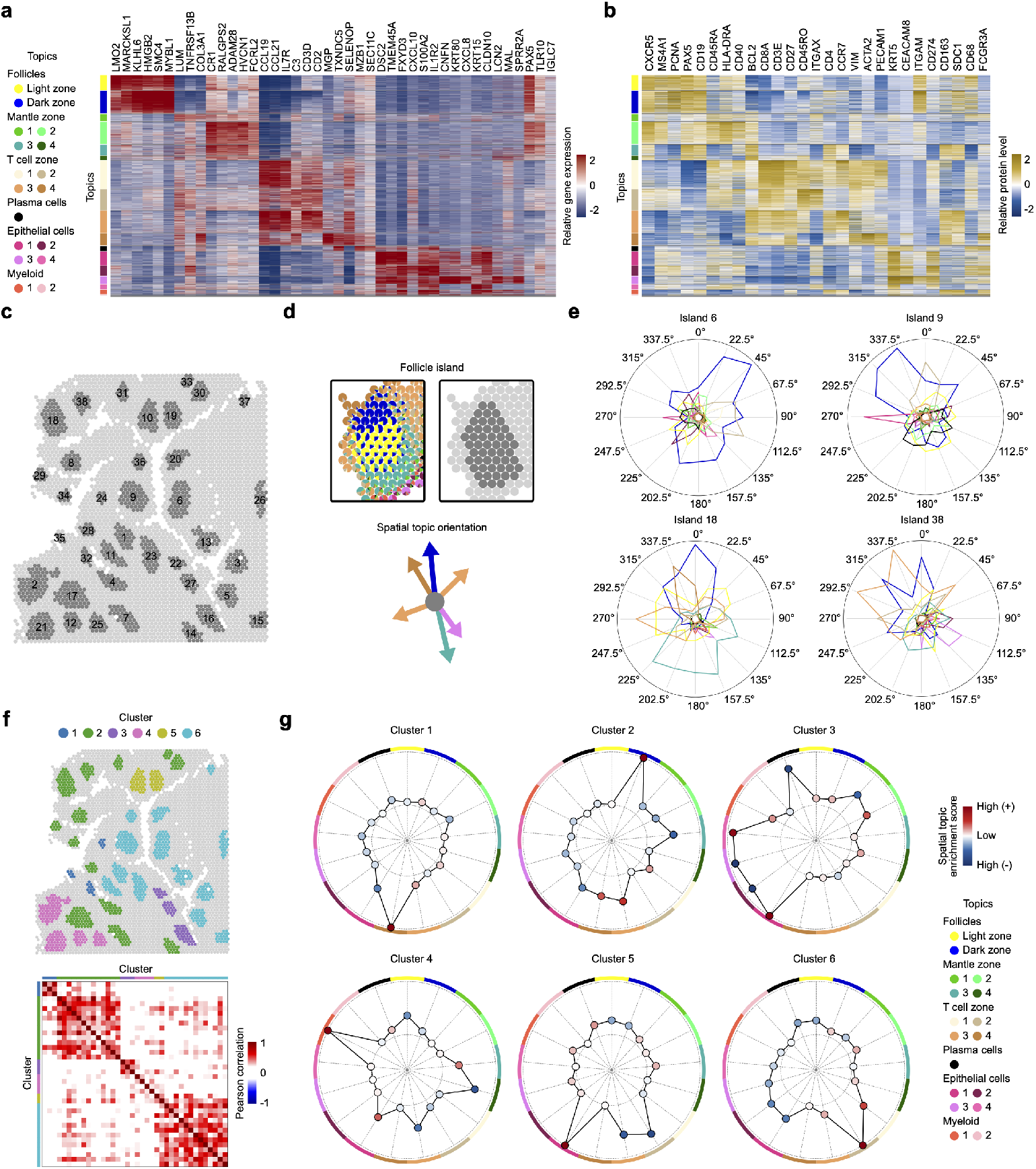
Extended analysis of the Human tonsil dataset. **a**, Heatmap of the top 3 RNA markers per topic. **b**, Heatmap of all protein markers in the panel. **c**, Spatial organization of follicle islands. **d**, Schematic illustrating interpretation of topic presence around island centroids. **e**, Radial profiles showing topic presence around centroids of selected islands. **f**, Clustering of follicle islands based on their radial profiles. **g**, Cluster-averaged radial profiles of follicle islands.

**Extended Data Fig. 4.**
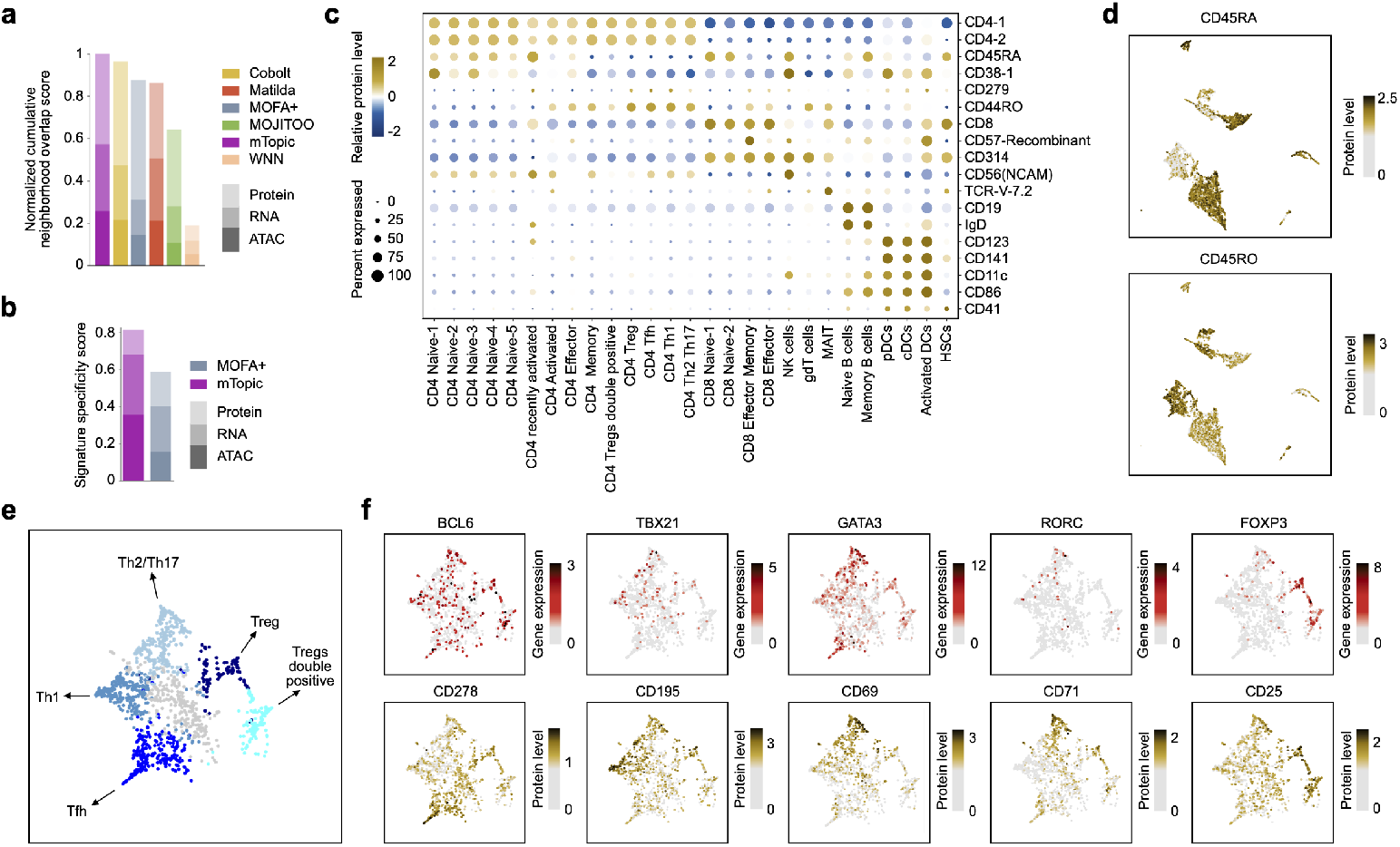
Benchmarking and identification of CD4 T cell subpopulations in the PBMC DOGMA-seq dataset. **a-b**, Benchmarking preservation of cell neighborhoods (a) and specificity of feature signatures (b) by various multimodal methods. **c**, Dotplot of protein markers per cell identity. **d**, Feature plots showing distribution of CD45RA and CD45RO markers that demarcate naive and effector T cells, respectively. **e**, Identification of CD4 effector T cell subsets. **f**, Expression of specific markers of CD4 T cell subpopulations, from the left: Tfh, Th1, Th2, Th17, and Treg. The upper panel shows RNA levels of specific TFs, and the bottom panel shows the levels of protein markers.

**Extended Data Fig. 5.**
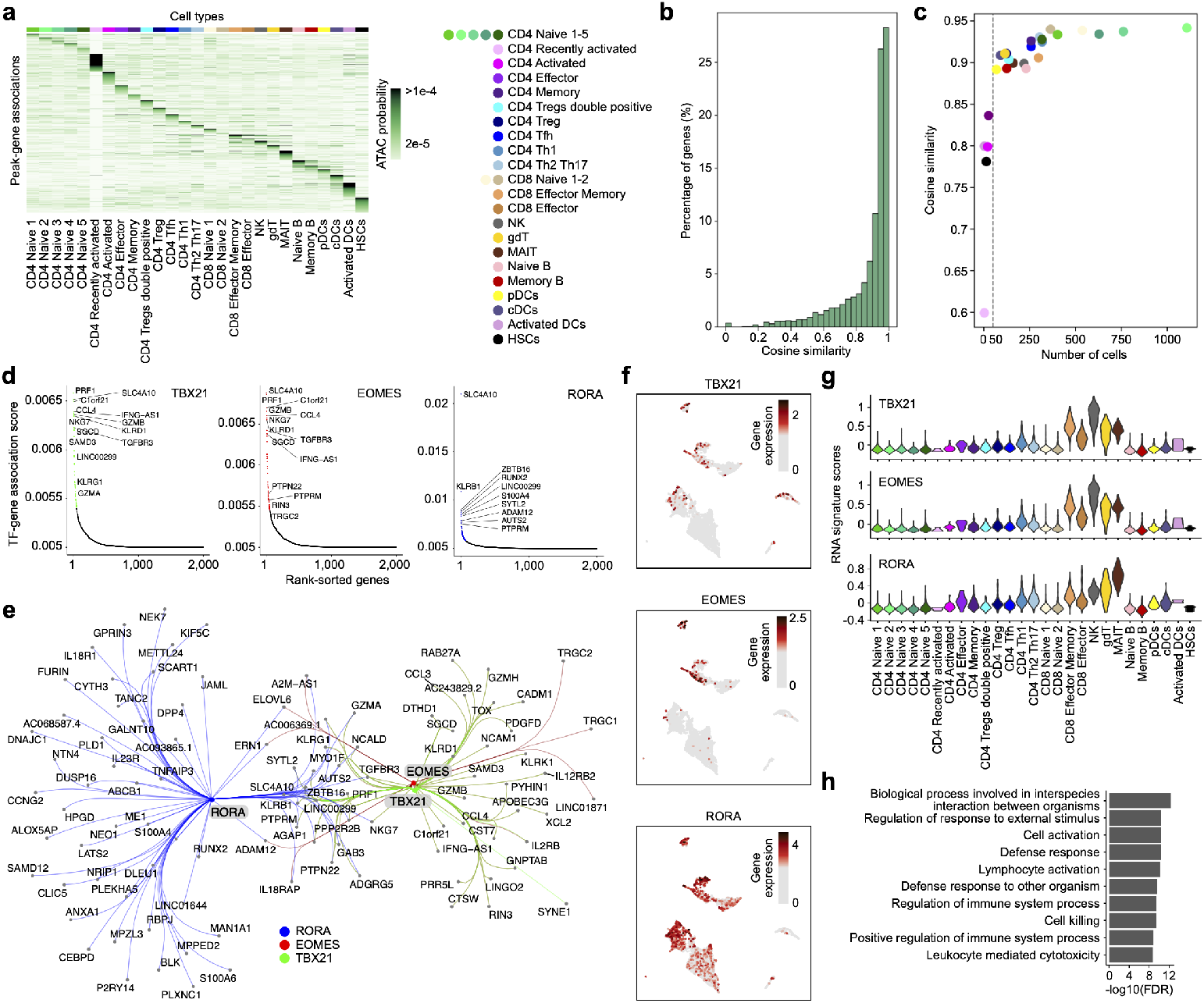
Peak-gene association characteristics (a-c) and extended TFs analysis for NK and cytotoxic T cell populations in PBMC DOGMA-seq dataset (d-h). a, ATAC scores across inferred peak-gene associations. **b-c**, Cosine similarity between gene expression inferred from peak-gene associations based on chromatin accessibility levels and observed gene expression levels. **b**, Distribution of the cosine similarities across all genes from (**a**), **c**, Dependence of the cosine similarity on the number of cells assigned to topics. **d**, Rank-sorted TF-gene associations for TBX21, EOMES, RORA. **e**, Visualization of the GRN formed by TBX21, EOMES, RORA. **f**, Expression values of TBX21, EOMES, and RORA visualized on UMAP. **g**, Gene signature scores of TF targets across cell types. **h**, Pathway analysis of the TF targets.

