## Supplementary Information for "Identifying multimodal molecular programs with mTopic"

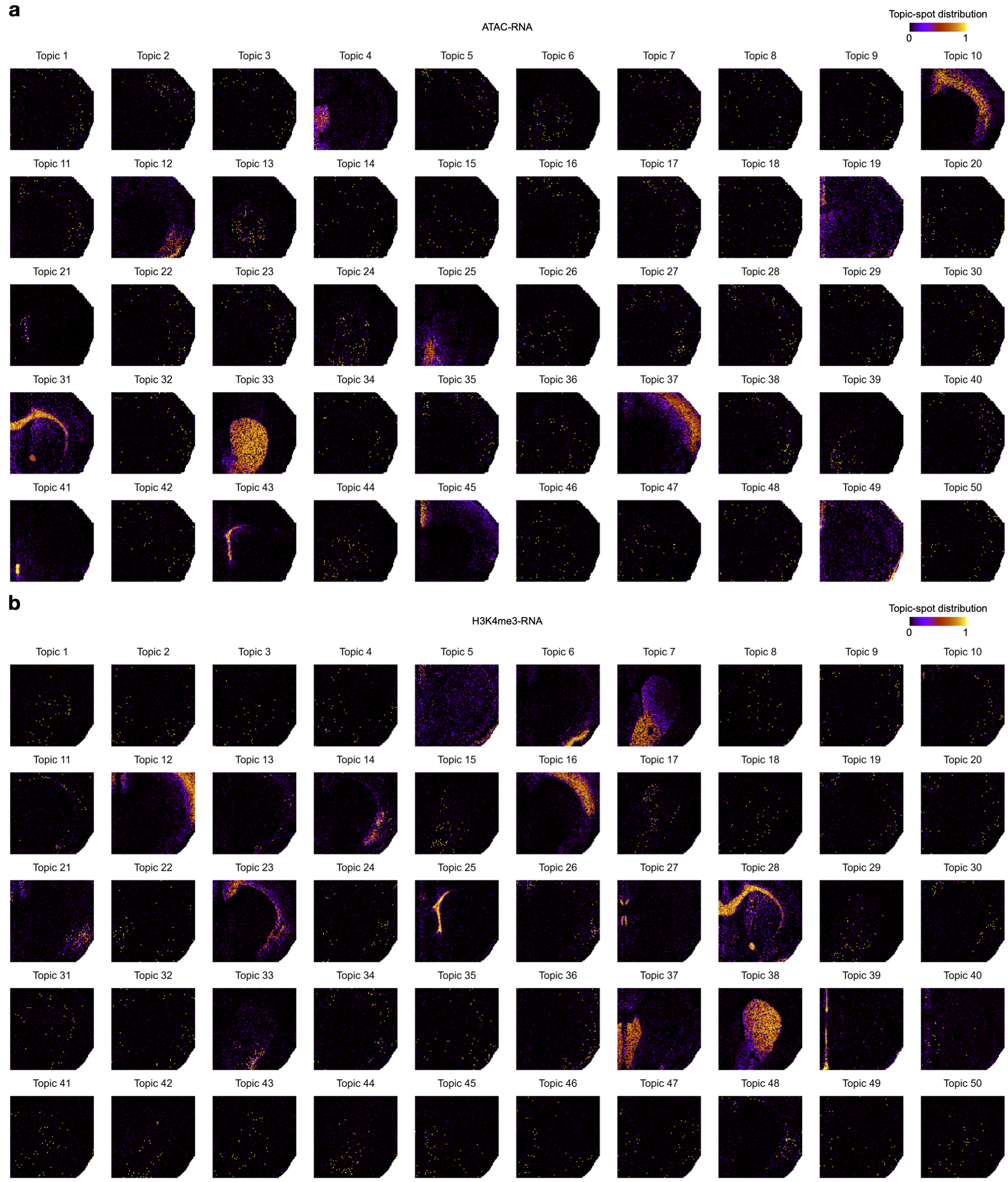


**Supplementary Fig. 1: Topic-spot distributions computed using mTopic for P22 mouse brain datasets** [1]**.** **a**, ATAC-RNA dataset. **b**, H3K4me3-RNA dataset.

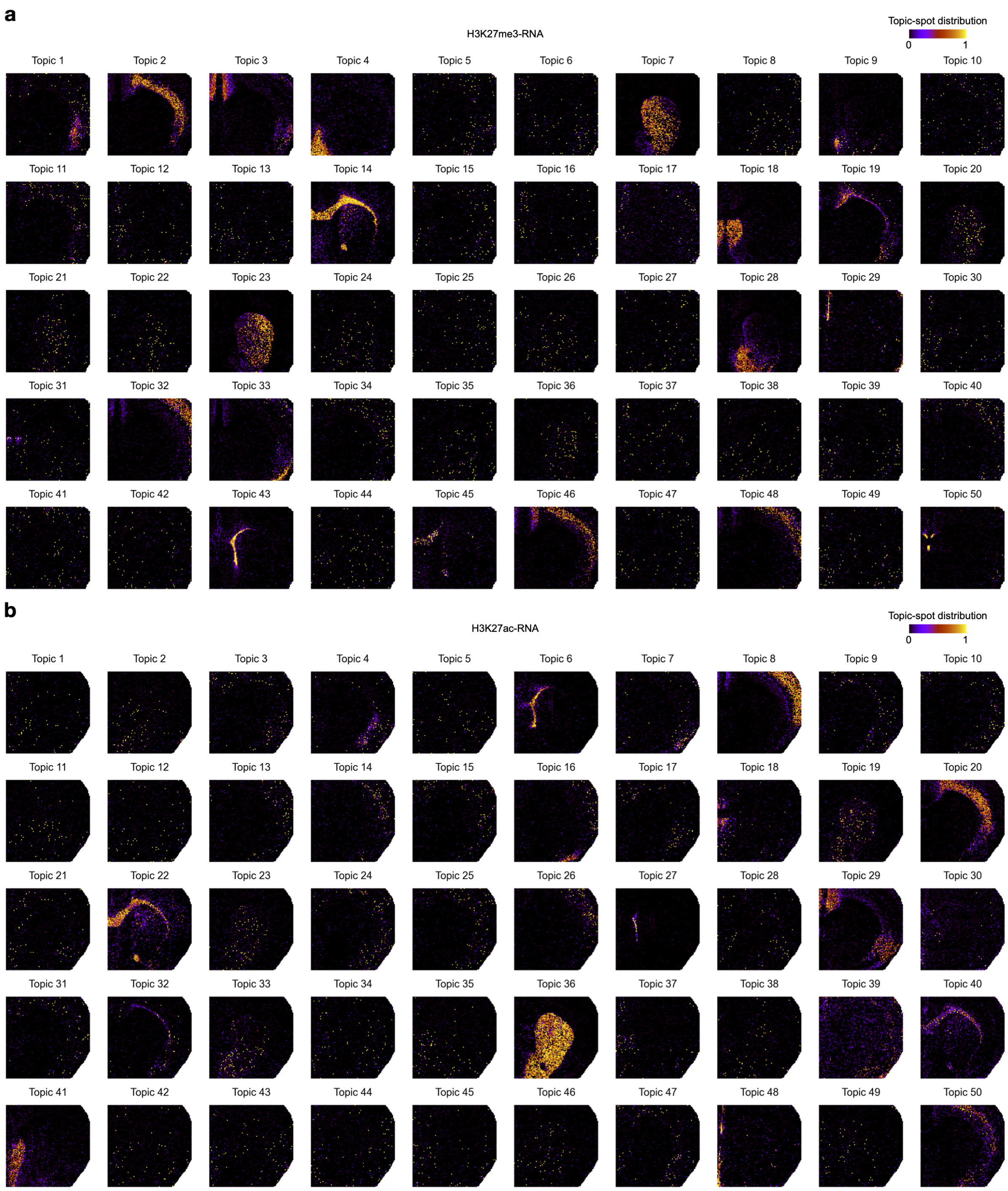


**Supplementary Fig. 2**: **Topic-spot distributions computed using mTopic for P22 mouse brain datasets [1]. a**, H3K27me3-RNA dataset. **b**, H3K27ac-RNA dataset.

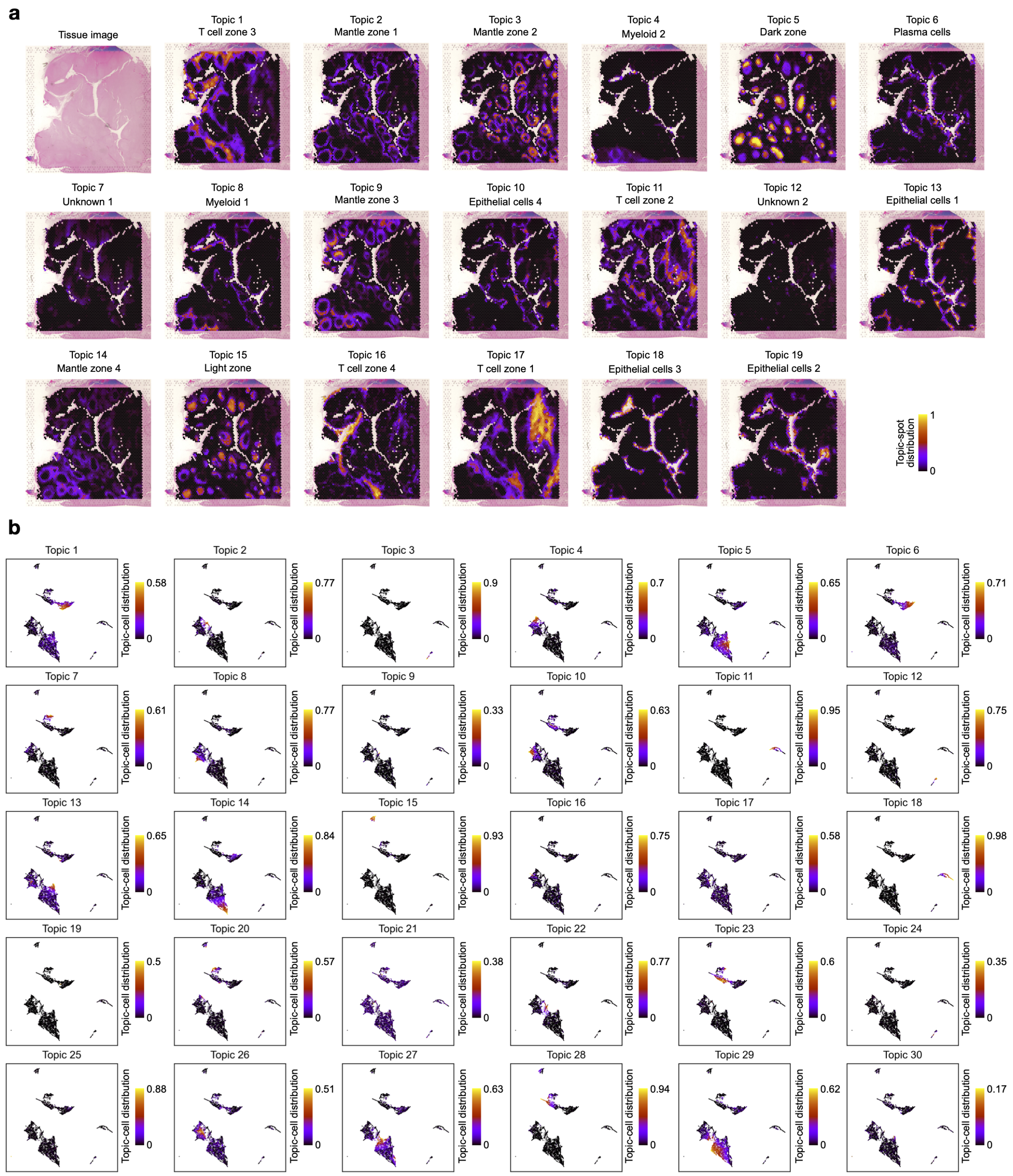


**Supplementary Fig. 3**: **Topic distributions computed using mTopic for human tonsil and PBMC datasets [2, 3].** **a**, RNA-Protein human tonsil dataset. **b**, ATAC-RNA-Protein PBMC dataset.
